# Quantitative modelling of signaling in aggressive B cell lymphoma unveils conserved core network

**DOI:** 10.1101/2024.03.25.586526

**Authors:** Bertram Klinger, Isabel Rausch, Anja Sieber, Helmut Kutz, Vanessa Kruse, Marieluise Kirchner, Philipp Mertins, Arnd Kieser, Nils Blüthgen, Dieter Kube

**Affiliations:** Institute of Pathology, Charité-Universitätsmedizin Berlin, corporate member of Freie Universität Berlin and Humboldt-Universität zu Berlin, Berlin, Germany; Institute for Theoretical Biology, Humboldt-Universität zu Berlin, Berlin, Germany; Clinic of Hematology and Medical Oncology, University Medical Centre Goettingen, Göttingen, Germany; ZytoVision GmbH, Bremerhaven, Germany; Research Unit Gene Vectors, Helmholtz Center Munich - German Research Center for Environmental Health, Munich, Germany; Core Unit Proteomics, Berlin Institute of Health at Charité - Universitaetsmedizin Berlin and Max-Delbrueck-Center for Molecular Medicine, Berlin, Germany; German Cancer Research Center (DKFZ), Heidelberg, Germany; German Cancer Consortium (DKTK), Partner Site Berlin, Berlin, Germany; Berlin Institute of Health (BIH), Berlin, Germany; Research Unit Signaling and Translation, Helmholtz Center Munich - German Research Center for Environmental Health, Neuherberg, Germany; German Center for Infection Research (DZIF), Partner Site Munich, Germany

**Keywords:** STASNet, lymphoma, oncogenic signaling, conserved B cell signaling network, negative crosstalk p38 -| MEK/ERK

## Abstract

B cell receptor (BCR) signaling is required for the survival and maturation of B cells and is deregulated in B cell lymphomas. While proximal BCR signaling is well studied, little is known about the crosstalk of downstream effector pathways, and a comprehensive quantitative network analysis of BCR signaling is missing. Here, we semi-quantitatively modelled BCR signaling in Burkitt lymphoma (BL) cells using systematically perturbed phosphorylation data of BL-2 and BL-41 cells. The models unveiled feedback and crosstalk structures in the BCR signaling network, including a negative crosstalk from p38 to MEK/ERK. The relevance of the crosstalk was verified for BCR and CD40 signaling in different BL cells and confirmed by global phosphoproteomics on ERK itself and known ERK target sites. Compared to the starting network, the trained network for BL-2 cells was better transferable to BL-41 cells. Moreover, the BL-2 network was also suited to model BCR signaling in Diffuse large B cell lymphoma cells lines with aberrant BCR signaling (HBL-1, OCI-LY3), indicating that BCR aberration does not cause a major downstream rewiring.

## Introduction

Intracellular signaling pathways are central to the communication of a cell with its environment, and control many important cellular processes and fates. These pathways are often activated by ligands that bind to cognate cell surface receptors and activate a specific set of intracellular proteins, often by tyrosine or serine/threonine phosphorylation. In many disease contexts, these pathways are deregulated, for instance by mutations in key signaling proteins. Consequently, activation can occur independent of ligands or stimuli. Signaling pathways are embedded into complex networks with feedback and crosstalk. It is therefore difficult to predict how these networks change when a pathway is chronically activated, and how the pathway reacts upon targeted interference. Mathematical modeling of intracellular pathways based on systematic perturbation data is a valuable approach to disentangle the difference in signaling networks of ligand/stimuli-induced pathways vs. chronic (aberrant) active pathways. A better understanding of the interactions of chronically activated signaling pathways is important to improve the prediction and further design of targeted therapies.

B cell receptor signaling represents a network for which the extend of feedback and crosstalk still remains unclear. Furthermore, a better dissection of chronic vs. acute signaling will support our understanding of related diseases. In normal B cells, the pathway is triggered by antigens, but is chronically activated in specific sets of Non-Hodgkin B cell lymphoma (NHL). In normal physiology the binding of a specific antigen to the B cell receptor complex leads to a very fast rebuilding and recruitment of a number of proximal signaling molecules including the ITAM motif molecules CD79A/B, the kinases LYN/SYK and SLP65/BTK/PI3K (Cantrell, 2015; Pal Singh et al., 2018; Schulze-Luehrmann & Ghosh, 2006). The activated BCR subsequently recruits multiple downstream signaling molecules and pathways, some of which include those dependent upon phospholipase Cγ, Protein kinase C or RAF-MAPK, PI3K, GSK3, MTOR and NF-kB. In addition, also JNK and p38MAPK are activated downstream of the BCR (Johnson & Lapadat, 2002). These pathways converge in the activation of a set of different transcription factors controlling B cell proliferation and survival, including c-Myc, NF-AT, Elk1, c-Jun and ATF2 (Efremov et al., 2020; Johnson & Lapadat, 2002; Satpathy et al., 2015).

In human B cells the activation of the key signaling pathway RAF/MEK/ERK by BCR seems to be similar to the activation by receptor tyrosine kinases (Satpathy et al., 2015; Vanshylla et al., 2018). This signal transduction network downstream of the B cell receptor (BCR) has received much attention, as it is a major regulator in adaptive immunity (Casola et al., 2019; Davis et al., 2010; Havranek et al., 2017; Schmitz et al., 2012). It has been shown that the relative activity of the multiple pathways downstream of BCR determines the outcome of BCR signaling (Reth & Brummer, 2004). The activity of the BCR is important for the expansion and survival and can be supported by both antigen-dependent and antigen-independent mechanisms (Efremov et al., 2020). Compelling evidence is provided that in different NHL subtypes aberrant BCR signaling activation is important (Davis et al., 2010; Elenitoba-Johnson & Lim, 2018; Lenz & Staudt, 2010; Pasqualucci & Dalla-Favera, 2018; Phelan et al., 2018; Schmitz et al., 2014; Shaffer et al., 2012; ten Hacken & Burger, 2016). BCR signaling is often studied in Burkitt lymphoma (BL) cell lines (Efremov et al., 2020; He et al., 2018a; Satpathy et al., 2015; Schrader et al., 2012; Vanshylla et al., 2018).

While the EGFR-signaling network or the T cell receptor pathway have been quantitatively studied in many contexts, the BCR signaling and its oncogenic variations have so far only gained limited attention. Elaborated network models of EGFR and T cell signaling include proximal and distal elements but also feedback loops (Cantrell, 2015; Chaudhri et al., 2010; Kholodenko, 2000; Klinger et al., 2013; Kumar et al., 2007; Saez-Rodriguez et al., 2007; Schoeberl et al., 2002; Young et al., 2019). Some of these models discovered and quantified pathway interactions and feedback loops, which have important implications also for clinical applications. These include positive feedback loops that strengthen the therapeutic success and negative feedback loops that lead to adverse events or even therapeutic failure (An et al., 2020; Gu et al., 2022; Klinger et al., 2013; Sun et al., 2014, Di Nicolantonio et al., 2021). For example, the somewhat disappointing response rates to drugs targeting the intracellular RAS/MAPK pathway could be explained by strong negative feedbacks (Klinger et al., 2013; Prahallad et al., 2012). Now it is becoming more and more clear that also in BCR signaling the signal is propagated through a complex context-dependent network. The complexity of this signaling network is also characterized by an interplay of RAF/MEK/ERK, JNK, p38, NF-kB and PI3K, originally described as parallel pathways of activation. However, there is a substantial crosstalk between these different downstream signaling molecules not yet characterized in sufficient detail. Reaching a better level of understanding will allow to answer questions such as whether these crosstalks are cell type specific and thus account in part for the differential activity of BCR and its functional outcome, but can also unveil therapeutic opportunities.

The few existing BCR signaling models are mostly deterministic models that include BCR signal propagation by a specific set of network nodes (Barua et al., 2012; Chaudhri et al., 2010; Kumar et al., 2007; Mcgee et al., 2015). The models cover both membrane proximal, early signaling events and to some extend downstream signaling events. Furthermore, a detailed model of the feedback loops involving LYN and SYK incorporating every phosphorylation event for six proximal signaling components has been established (Barua et al., 2012). However, an exhaustive quantitative modelling of downstream pathways is currently missing.

In a previous study Pirkl et al., 2016 attempted to infer an oncogenic signaling network in aggressive lymphoma cell lines indirectly from gene expression data following signaling perturbations using the Boolean Nested Effects Modelling (B-NEM) approach. By this simple modelling approach, an acyclic network with only activating interactions was derived which contained no feedbacks and lacked quantitative information. While such models help to gain the understanding of pathway interactions, they are not suitable for simulations or quantitative model comparisons. To get a more fine-grained semi-quantitative understanding of the signaling network, we therefore decided to employ a modelling approach on the more information rich readout of phosphorylation data after systematic pathway perturbations which contains more predictive power (Eduati et al., 2017). We have previously developed an approach termed STASNet that is based on Modular Response Analysis (MRA) (Kholodenko et al., 2002) and applied this tool to decipher EGFR/RAS signaling in different tumors (Dorel et al., 2018; Hood et al., 2019; Klinger et al., 2013) as well as to compare mouse embryonic stem cells with different sex chromosome compositions (Sultana et al., 2023). This approach would allow to model and identify feedback loops that are known to play a crucial role in conferring robustness, shaping the signaling dynamics or integrating multiple converging signals and thus are vital to consider when searching for drug combinations eligible for targeted therapeutics. Furthermore, in contrast to the previous Boolean approach, STASNet allows to (semi)-quantitatively compare signaling networks and to predict simulations across cell line models.

Based on systematic perturbation data we developed an MRA-based network model for acute signaling in a Burkitt lymphoma (BL) cell line BL-2. We show that this model is transferable to perturbation data of another BL cell line BL-41. This supports the view that our model reflects common elements of intracellular signaling of acute B-cell signaling. Additional phosphoproteomic analyses of selected interventions support key insights of this model. After establishing these quantitative network models, we found that the BL-2-derived network structure forms a solid base to describe signaling networks in cell lines from Diffuse large B cell lymphoma (DLBCL) with aberrant BCR signaling. Thus, by integrated phosphoproteomic analysis, we have uncovered a conserved core B cell receptor-regulated signaling network.

## Material and Methods

### Cell culture and stimulation

BL-2, BL-41, and OCI-Ly3 cells were obtained from the DSMZ (Braunschweig, Germany). HBL-1 cells were kindly provided by D. Krappmann (Munich, Germany). BL-2 and BL-41 cells were cultivated as described previously at cell densities between 2×10^5^ and 1×10^6^ cells/ml (Vockerodt et al., 2005). For stimulation studies, BL-2 and BL-41 cells were cultured in RPMI1640 with 10 % FCS at 3×10^5^ cells/ml and incubated with 1.3µg/ml goat α-IgM F(ab)2 fragments (Jackson Immunity) for indicated time points. For pathway interventions cells were treated with DMSO, 5µM SP600125, (all Merck KGaA, Darmstadt, DE), 1µM AZD6244, 2µM BKM120, 10µM Ibrutinib, 1µM MK-2206, 1µM Rapamycin (all Selleckchem, Munich, DE), 10µM MLN120B (MedChemExpress, Sollentuna, SE), 2µM SB203580 (Sigma-Aldrich, St. Louis, US). HBL-1 or OCI-LY3 cells were cultured in RPMI1640 with 10 % FCS at 5×10^5^ cells/ml. For studying protein phosphorylation, the cells were incubated with inhibitors for 3 hours. HBL-1 and OCI-LY3 were then harvested, while BL-2 and BL-41 cells were incubated for additional 30 minutes with 1.3µg/ml goat α-IgM F(ab)2 fragments (BCR activation) or left untreated. Cells were harvested using corresponding inhibitors of phosphatases (PhosSTOP Roche, Basel, CH) and proteases (cOmplete Mini Roche, Basel, CH) and protein was isolated using RIPA buffer.

### Western blot analysis

Cells were analysed for protein expression by SDS polyacrylamide gel electrophoresis and Western blot analysis using the following antibodies: mouse monoclonal anti α-tubulin (#05-829, Merck Millipore, Burlington, US), rabbit α-p-AKT^Ser473^ (#9271), rabbit α-pan AKT (#9272), rabbit α-p-p38 (#9211), rabbit α-p38 (#9212), rabbit α-p42/44 (#4695), mouse α-MEK1/2 (#4694), rabbit α-p-p42/44 (#4370), rabbit α-p-p42/44 (#4377), rabbit α-p-MEK1/2 (#9154), rabbit α-p70 S6 Kinase (49D7) (#5707), rabbit α-p-p70 S6 Kinase^Thr389^ (108D2)(#9234), α-p-Raf1^Ser289/296/301^(#9431), rabbit α-p-Raf1^Ser338^ (56A6) (#9427)(all from Cell Signaling Technology) and α-mouse HRP polyclonal goat (D1609) and α-rabbit HRP polyclonal goat (E1710) (all from Santa Cruz Biotechnology, Inc.).

### Bio-Plex^®^ multiplex immunoassay

To measure all protein phosphorylations in one sample, the magnetic bead-based multiplex assay (BIO-RAD) was performed. The principle is that a specific antibody coupled to a color-coded bead identifies the total protein of interest while a second detection antibody determines the magnitude of a distinct phosphorylation. For this analysis, the Bio-Plex Pro Cell Signaling Reagent Kit (BIO-RAD) was used. The treated lymphoma cells were cooled down by addition of the three-fold volume of ice-cold DPBS supplemented with 1xPhosSTOP (Roche) and 100µM sodium orthovanadate. After 5 minutes centrifugation (500xg, 4°C) the cells were washed once. According to the instruction manual the cells were lysed in the provided buffer containing 1xfactor QG and 2mM PMSF. After shaking for 20 minutes at 4°C, debris was removed by centrifugation (14000xg, 4°C) for 15 minutes. The analysis with the Bio-Plex Protein Array system (BIO-RAD) was done as published before (Klinger et al., 2013) and according to the manufacturer’s instructions. Specific beads were used for p-SYK^Y352^, p-ZAP70^Y319^, p-BTK^Y223^, p-AKT^S473^, p-40S ribosomal protein S6^S235/S236^, p-BAD^S136^, p-MEK1^S217/S221^, p-ERK1/2^T202/Y204, T185/Y187^, p-p90RSK^S380^, p-GSK3αß^S21/S9^, p-HSP27^S78^, p-JNK^T183/Y185^, p-c-Jun^S63^ and p-p65 NF-κB^S536^. The Bio-Plex manager software and R package lxb was used for data acquisition.

### Network modelling

For the quantitative network modeling from systematic perturbation data we used our previously developed R package STASNet Version 1.0.2 (available on https://github.com/molsysbio/STASNet), which a derivative of Modular Response Analysis (Bruggeman et al., 2002; Kholodenko et al., 2002) adapted to model incomplete signaling perturbation data (Dorel et al., 2018; Klinger et al., 2013). The modelling strategy is illustrated in **FIGURE 1B** and accompanying text. More detailed modelling workflows as well as the required data for cell lines BL-2, BL-41, HBL-1 and OCI-LY3 can be found in html reports stored in zenodo (https://zenodo.org/doi/10.5281/zenodo.10732059).

**FIGURE 1.**
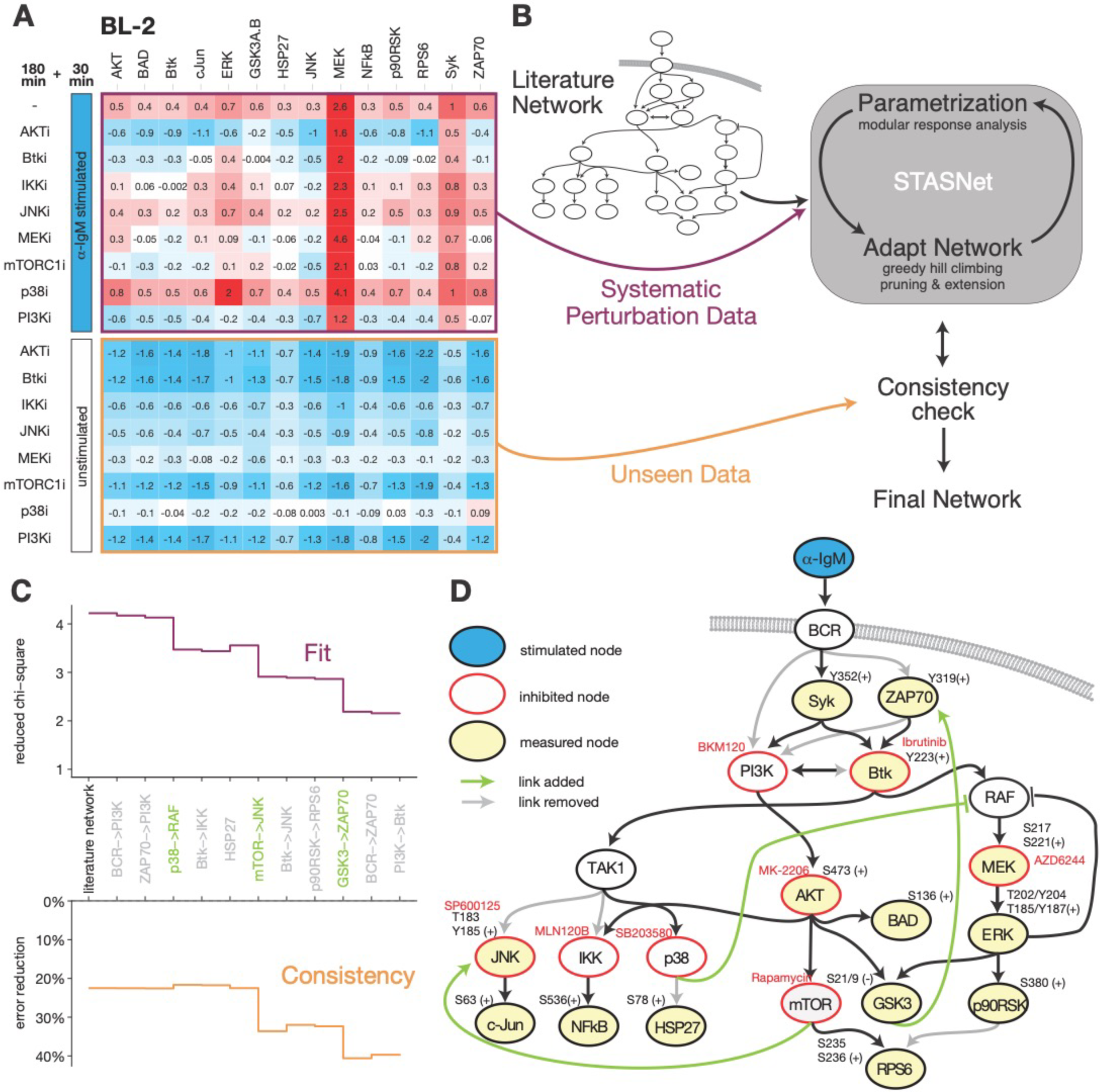
Discovery of novel feedback and crosstalk structures in a BCR-driven signaling network by perturbation data-based modelling of BL-2 cells. (A) Systematic perturbation data shown as log2 fold changes to solvent control DMSO. Data was generated by pre-treating BL-2 cells with inhibitors targeting key effectors downstream of BCR for 3h with subsequent BCR stimulation using α-IgM for 30 min (upper panel) or no stimulation as consistency check (lower panel). Phosphorylation of indicated signaling proteins (cf. **D**) was measured using bead-based ELISAs (n=3). (B) Modelling workflow using the Modular Response Analysis-based method STASNet to derive a semi-quantitative directed network. The model requires systematic perturbation data (depicted in **A**) and a curated literature network as starting network (cf. **D**). In order to avoid overfitting, the data was split into two parts: (1) α-IgM stimulated data was used for parameter fitting and network adjustment and (2) unstimulated inhibitor data was used for verifying model consistency. After each network adjustment step the unseen data part was simulated and compared. If the error reduction as compared to the null model was not significantly worse, the new network adjustment was upheld otherwise the next best solution was simulated and tested. (C) Model performance for each modelling step from the literature-derived starting model to the final model: (TOP) goodness of fit as weighted sum squared residuals divided by number of free parameters and (BOTTOM) consistency check step as percentage of error reduction compared to unperturbed control as null model. (D) Literature network adjusted to the final signaling network for BL-2 cells derived by the modelling pipeline depicted in **B**. grey line/text - removed links/nodes; green line/text - added links.

In general, to assess goodness of fit we use the sum weighted squared residuals (WSSR) which is defined as:

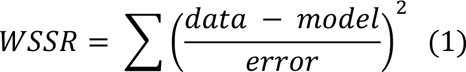

where the data represents the data points, the model the prediction and error the antibody-wise standard error derived from the replicate measurements. Using equation (1) for the model prediction of the fitted data (WSSRfit) we calculated the reduced chi square statistics Xr as goodness of fit assessment, which is defined as:

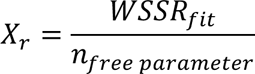

Where the number of free parameters is given by number of data points minus the number of model parameters. For the model consistency step the STASNet function *simulateModel()* was used to generate the prediction of unstimulated but inhibited data of which the sum of weighted squared residuals (WSSRsimulation) was calculated. We then compared this to the residuals that a suitable null model, i.e. unperturbed data, would produce (WSSRnullmodel) and defined the prediction capacity better than null model as follows:

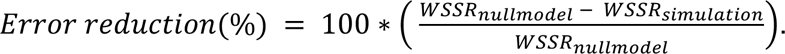

For comparative combined modelling we used the following strategy. Since inhibitor dosages were always the same but the response to the inhibitors varied from cell line to cell line, the actual difference is thought to happen on the internal wiring not on the inhibitor perturbation strength parameter. Therefore, when comparing different models, we fixed the parameters encoding the inhibitor strengths to the same value across the models to be compared and allowed the STASNet algorithm to compensate by refitting the remaining parameters using the STASNet routine *refitModel()*.

### Mass spectrometry based phosphoproteomics

For phosphoproteomic profiling by mass spectrometry, cell pellets were lysed in urea lysis buffer (6M urea, 2M thiourea, 100mM Tris-HCl, pH 8, 150mM NaCl, 1mM EDTA, phosphatase inhibitor cocktail 2 and 3, 10mM NaF; Sigma), reduced with 10mM DTT (dithiothreitol, Sigma) for 45 minutes followed by alkylation with 40mM CAA (2-chloroacetamide, Sigma) for 30 minutes. After treatment with Benzonase® (Merck, 50 units) for 30 min at 37°C, samples were centrifuged for 10 minutes at 12 000 rpm. The supernatant was collected and protein concentration was determined. 200µg protein per sample was digested with 2µg endopeptidase LysC (Wako), followed by a 3:1 dilution with 100mM ammonium bicarbonate and addition of 2µg sequence-grade trypsin (Promega). Samples were digested at room temperature overnight and acidified with formic acid (final concentration 1%). The resulting peptides were cleaned using C18 SepPak columns (Waters, 100mg/1cc), dried and resolved in 50mM HEPES (pH 8). Peptides were labeled with 11-plex tandem mass tag (TMT, Fisher Scientific) reagents following the vendors instructions. After combining all samples and C18 SepPak-based clean-up (Waters, 200mg/1cc), samples were fractionated by high-pH reversed phase off-line chromatography (1290 Infinity, Agilent) and pooled into 15 fractions, which were applied to IMAC based phosphopeptide enrichment as described (Mertins et al., 2018). For LC-MS/MS measurements, peptides were reconstituted in 3% acetonitrile with 0.1% formic acid and separated on a reversed-phase column (20 cm fritless silica microcolumns with an inner diameter of 75 µm, packed with ReproSil-Pur C18-AQ 1.9 µm resin (Dr. Maisch GmbH)) using a 98 min gradient with a 250 nl/min flow rate of increasing Buffer B (90% ACN, 0.1% FA) concentration (from 2% to 60%) on a High Performance Liquid Chromatography (HPLC) system (Thermo Fisher Scientific) and analyzed on a Q Exactive Plus instrument (Thermo Fisher Scientific). The mass spectrometer was operated in data-dependent acquisition mode using the following settings: full-scan automatic gain control (AGC) target 3 x 10^6^ at 70K resolution; scan range 350–2000 m/z; Orbitrap full-scan maximum injection time 10ms; MS/MS scan AGC target of 5 x 10^4^ at 35K resolution; maximum injection time 100ms; normalized collision energy of 32 and dynamic exclusion time of 30s; precursor charge state 2-6, ten MS2 scans per full scan. RAW data were analyzed with MaxQuant software package (v 1.6.0.1) using the Uniprot databases for human (2018-05). The search included variable modifications of methionine oxidation, N-terminal acetylation, deamidation (N and Q) and phosphorylation (STY) and fixed modification of carbamidomethylated cysteine. Reporter ion MS2 for TMT11 was selected (internal and N-terminal) and TMT batch specific corrections factors were specified. The FDR (false discovery rate) was set to 1% for peptide and protein identifications. Unique and razor peptides were included for quantification.

Data availability: Mass spectrometry raw files and processed MaxQuant output files are available on the PRIDE proteomics database with identifier PXD047709.

### Statistical analysis and evaluation of Tandem Mass Tag data

After excluding reverse database hits and potential contaminants the resulting list of phosphosites was filtered for localisation probability (>0.75) and unified for ambiguously mapped phosphosites leaving 28871 phosphosites for downstream analysis. After quantile normalisation, differentially regulated sites were identified using the limma R package (v. 3.54.0) and resulting p-values were corrected for multiple testing using Benjamini-Hochberg method (FDR<=0.05).

Kinase enrichment of phosphopeptides was conducted by using information from PhosphoSitePlus (https://www.phosphosite.org, Kinase_Substrate_Dataset 24.11.2021) to identify putative kinases that regulate the phosphosites. Then for all kinases that regulate at least 5 phosphosites in our data set, we tested with a two-sided paired t-test whether the mean fold change to the reference treatment (i.e., Control+DMSO for α-IgM+DMSO and α-IgM+DMSO for α-IgM+inhibitor) was significantly different from 0. Afterwards we corrected for multiple-testing using the Benjamini-Hochberg approach. To assess the effect of treatments on the total proteome we in parallel analyzed the total protein data (n=8785) analogously to the phosphoproteome data but without localization probability filter. In total only 71 proteins were found significantly changed for any treatment of which 35 overlapped with significant phosphosite changes, affecting 1.1% of proteins whose phosphoproteome significantly changed and 1.6% of significant phospho-signals (n=141). Due to the little change of total proteins we decided to only concentrate on the phosphoproteome in this study.

## Results and Discussion

To dissect B cell signaling in aggressive NHL cells and get better insights into corresponding oncogenic signaling networks, we followed an approach that we previously established for EGFR signal transduction networks in solid cancer cell lines (Dorel et al., 2018; Klinger et al., 2013). This approach combines experimental quantitative perturbation data sets with a semi-mechanistic computational modelling approach derived from MRA.

### Generation of signaling perturbation data set

We decided to generate the first model on the Burkitt lymphoma cell line BL-2 as this type of lymphoma is a role model for studying acute B cell signaling (Denépoux et al., 2000; He et al., 2018b). In BL-2 cells an external stimulus (α-IgM) is able to activate the BCR and its downstream intracellular signaling (Denépoux et al., 2000; Poltoratsky et al., 2001; Schrader et al., 2012). Stimulation-mediated intracellular signaling typically displays a strong transient response with different kinetics followed by short or longer lasting activation plateau interval. Therefore, time-series experiments were performed to determine the optimal time point for the pathway intervention experiments. A strong response for AKT, ERK1/2, p70S6K and p38/MAPK14 over time in α-IgM treated BL-2 cells is observed using both immunoblotting and a bead-based ELISA platform (**Suppl. Figure 1**). The 30-min time point was chosen for further experiments, as the interpretation of the modeling procedure requires the signaling network to be approximately in steady state.

To establish an information rich dataset for modelling, we combined stimulation of the receptor with inhibitions of several key signaling nodes with targeted small-molecule inhibitors. Specifically, we preincubated inhibitors for JNK (5µM SP600125), MEK (1µM AZD6244), PI3K (2µM BKM120), BTK (10µM Ibrutinib), AKT (1µM MK-2206), MTOR (1µM Rapamycin), IKK (10µM MLN120B) and p38 (2µM SB203580) or solvent control for 3h. We then stimulated BCL-2 cells with α-IgM for 30min or left unstimulated (control). Before and after perturbation, phosphorylation of key signaling molecules was quantified to create a high-dimensional data set for a semi-quantitative database. A bead-based multiplex ELISA platform (MagPix) was then used to measure the phosphorylation of fourteen key signaling and effector proteins: SYK^Y352^, ZAP70^Y319^, BTK^Y223^, AKT^S473^, RPS6^S235/236^, BAD^S136^, ERK2^T185/Y187^, MEK1^S217/S221^, p90RSK^S380^, GSK-3a/b^S21/S9^, NF-κB-p65^S536^, HSP27^S78^, JNK^T183/Y185^ and cJun^S63^. The resulting data sets are presented as a heat map of log2 fold changes compared to unperturbed and unstimulated controls (**FIGURE 1A**).

### Model-based analysis of B cell receptor signaling in BL-2 cells

We used the perturbation data set together with a literature-based starting network (Efremov et al., 2020; Hendriks et al., 2014; Hood et al., 2019; Pal Singh et al., 2018) for our modelling pipeline to reverse engineer the signaling network downstream of the BCR (**FIGURE 1B and D**). We performed modeling using STASNet, a Modular Response Analysis-based modelling R package, to estimate optimal parameters for a given network structure to best match the perturbation data [Parametrization]. After fitting the initial network, we adapted the network structure to the specific cell system by removing links that did not contribute significantly to the goodness of fit (likelihood ratio test, p>0.05) and tested systematically if adding a link significantly improves the network fit (likelihood ratio test, p<0.05) [Adapt Network]. To prevent overfitting, structurally altered network models were validated by comparing simulations to data that were not used for model parameterization [Consistency]. These three modeling steps were iteratively repeated for every network change until no further alteration was supported by the data [Final Network]. **FIGURE 1C** shows the goodness of fit and the statistics of the consistency check during the network modeling steps. During the course of modelling the reduced chi-square statistics, representing goodness of fit, steadily decreases indicating better or equally goof fit with every network alteration. Next to the model fit a simultaneous consistency check procedure is conducted assessing the ability to model unseen data as percent reduction of weighted sum squared residuals (WSSR), when compared to untreated data as null model. For each modelling step the model predicts unseen data better or equally well than for the previous step. This is a good indication that the model development did not lead to overfitting.

**FIGURE 1D** presents the literature-derived starting network adjusted to the structure of the final BCR signaling network model for BL-2 cells with pruned links indicated in grey and novel links indicated in green. Four removed links correspond to redundancies in the receptor proximal signaling structure around PI3K, and removal resolved this non-identifiability. Other removed links belong to downstream signaling, decoupling JNK and IKK from BTK. Furthermore, RPS6 seems to be only regulated by AKT/mTOR and not ERK. HSP27 was not found to be regulated by p38 activity in BL-2 cells. Importantly, by extending the network, the modeling procedure discovered three novel connections that significantly improved the model fit: (i) a positive crosstalk from mTOR to JNK, (ii) a positive feedback from GSK3 to ZAP70 and (iii) a negative crosstalk from p38 on or above RAF/MEK/ERK.

### The network structure learned from BL-2 can be transferred to BL-41 perturbation data

After we have established a semi-quantitative network model of BCR signaling in BL-2 cells, we aimed to assess if the final network describes BCR signaling in a general way. Thus, we performed the same perturbations experiments that were conducted on BL-2 cells on a different BL cell line, BL-41, (**FIGURE 2A**) and subsequently applied the same modelling strategy (cf. **FIGURE 1B**) to derive a BL-41 specific signaling network. Starting from the same literature network as in BL-2 cells the network was derived that was best supported by the data. We noted that the reduced chi-square statistics approached the theoretical limit of 1 during network adaptation, indicating accurate model optimization (Fit, **FIGURE 2B**). Furthermore, the simultaneous independent consistency step (percentage of error reduction compared to unperturbed control) shows that the simulation quality of unseen data stayed approximately at a similar level during network development (Consistency, **FIGURE 2B**), indicating that no considerable overfitting took place.

**FIGURE 2.**
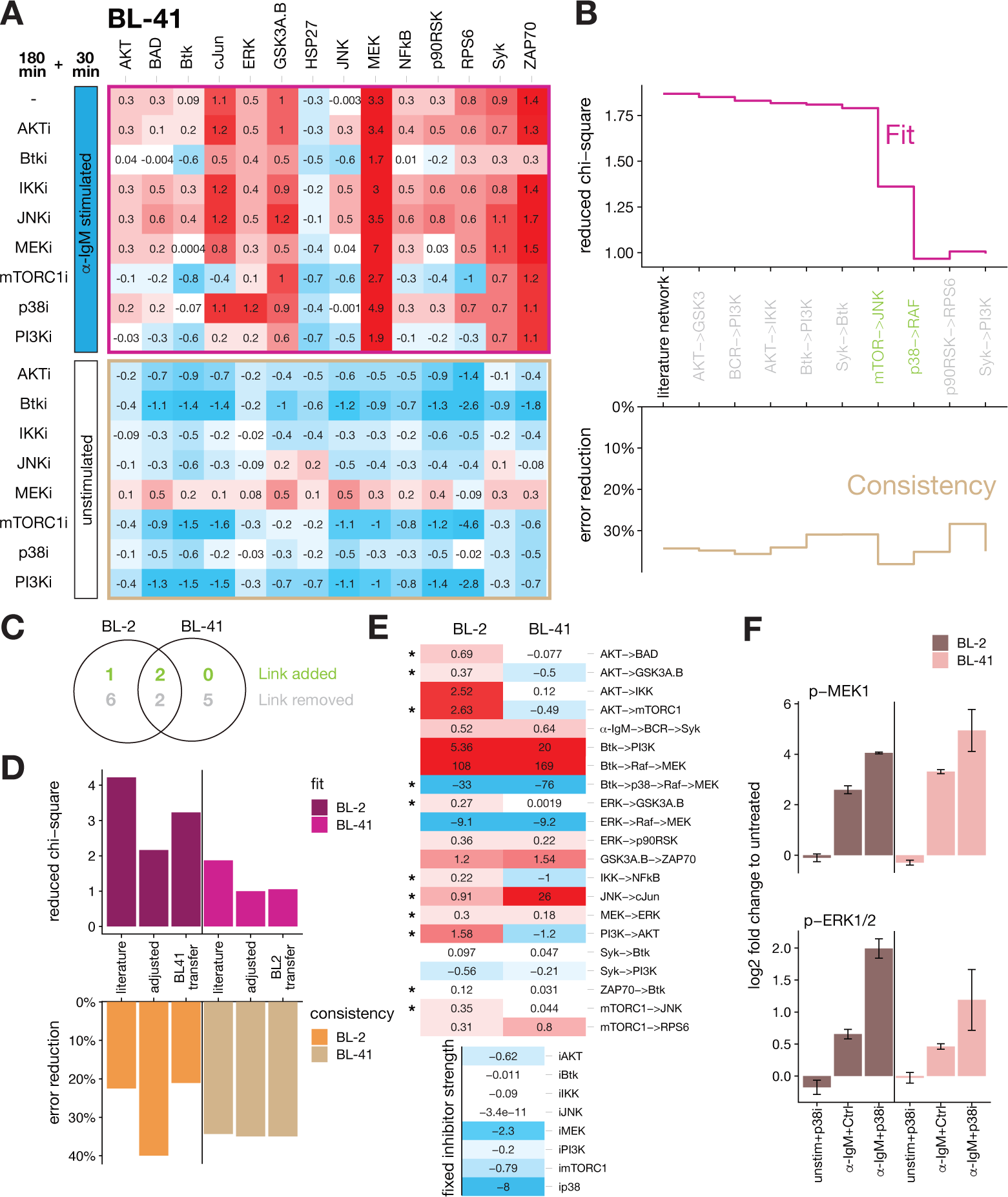
BL-2-derived modelling structure can be transferred to cell line BL-41. (A) Systematic perturbation data for BL-41 cells generated alike the procedure described in **FIGURE 1A**. (B) Model development statistics (TOP) goodness of fit as reduced chi-square and (BOTTOM) unseen data consistency check as percentage of error reduction compared to unperturbed control as null model for each modelling step from the literature-derived starting model (black and grey arrows in **FIGURE 1D**) to the final model (grey – reduction, green - extension). Modeling workflow see materials and methods. (C) Venn diagram indicating the shared and not shared structural adjustments in the development of BL-2 and BL-41 cells starting from the same literature network. (D) Model fit and consistency check statistics for fitted models on BL-2 and BL-41 perturbation data for three different network structures: (1) literature, (2) cell-specific adjusted network (adjusted) and (3) for the best-found structure of the respective other cell line (transfer). BL-2 structure is better transferable to BL-41 than *vice versa*. (E) Network coefficients heatmap from models fitted to the BL-2 learned structure for the indicated cell lines. Comparability was ensured by fixing the inhibitor strengths to BL-2-learned values as both cells received the same inhibitor doses. Stars denote coefficients that are significantly different (i.e., 95%-point wise confidence intervals do not overlap, see **Suppl. Table 1**). (F) Data excerpt for the model-derived negative crosstalk prediction from p38 to RAF/MEK/ERK pathway in BL-2 and BL-41 cells showing the upregulation of α-IgM-induced activation of pERK and pMEK by the p38 inhibitor SB203580 (mean ± s.e.m., n = 3), but no upregulation by p38 inhibitor alone.

During model adaption, 4 adjustments were shared between BL-2- and BL-41. This includes, two of the three additional links already identified in BL-2 cells before, i.e., p38->RAF and mTORC1->JNK. However, 7 and 5 adjustments were BL-2- and BL-41-specific, respectively (**FIGURE 2C**). This divergence in proposed network structure also manifests in the fact that only 14 of the 21 identifiable path coefficients were shared in both cell lines. A structural overlap of only 66% for cell lines from the same lymphoma type would indicate vastly different signaling. However, as our modelling technique employs a greedy hill climbing link adjustment strategy, the resulting networks may represent a local optimum. To further investigate this, we compared goodness of fit and model prediction (consistency) when the respective model structure of one cell line would be used to fit the data of the other cell line. **FIGURE 2D** shows the best fit and consistency statistics for the transferred models (transfer) in comparison to the literature-based network (literature) and the adjusted literature-based network (adjusted). Interestingly, the signaling network structure learned on BL-2 cells can be faithfully transferred to fit a BL-41 cell model with strikingly similar statistical properties than the best-found structure found for BL-41 itself. The reverse scenario, i.e., transfer of BL-41 best network structure to BL-2, led to a worse fit and consistency, while only slightly better than the literature network-based model.

Thus, it is evident that the current best network structure found for BL-2 cells can be transferred to BL-41 and results in an equally good model fit as if using the individually fitted network structure from BL-41 data. This successful transfer strongly supports the view of the similarity of BCR signaling in both BL cell lines. Furthermore, it indicates that the BCR signaling network structure developed from BL-2 data can be seen as the most representative BCR signaling structure for BL cells activated by a crosslink of the BCR (**FIGURE 1C**).

Next, we compared the fitted pathway coefficients of BL-2 cells and BL-41 cells using the BL-2-optimised network structure (**FIGURE 2E**). To improve comparability, the inhibitor strength parameters were fixed to values learned from the BL-2 model as both cell lines received the same doses of inhibitors. Thus, any change in the parameters should be attributed to intracellular signaling differences and not to differences in inhibitor strength. We performed a profile likelihood analysis on that unveiled that about half of the pathway coefficients (11 of 21, asterisks **FIGURE 2E**) are significantly different, i.e., the 95% pointwise confidence intervals (Raue et al., 2009) do not overlap. For example, the parameter corresponding to the pathway from Btk via p38 to MEK is estimated to be stronger in BL-41 than in BL-2 cells. Interestingly, this parameter corresponds to the cross talk from p38 to RAF/MEK/ERK, and the underlying data shows that the effect is only notable in the presence of α-IgM stimulus but not when p38 inhibitor is applied without stimulation (**FIGURE 2F**).

In summary, the BL-2 model constitutes a generic network structure that can accurately describe signaling for both BL cell lines. This finding is important as it demonstrates the existence of a signaling network core for the activated BCR in BL cells. In addition, three hitherto undescribed links in BCR mediated intracellular signaling in BL cells were unveiled to disentangle upstream wiring including the yet not fully resolved receptor proximal signaling events (**FIGURE 1C**). These so far undescribed network links include the strong negative crosstalk from p38 to RAF/MEK/ERK, which has been previously described in endothelial cells (Molinar-Inglis et al., 2022) or EGFR-activated epithelial cells (Finch et al., 2012). For BCR activated signaling this feedback however was not yet described. Therefore, we decided to conduct further experiments to characterize the model-predicted p38-mediated dampening of MEK1/2 phosphorylation.

### An α-IgM dependent feedback signal from p38 (MAPK14) attenuates pMEK and pERK

Since the model-based findings rely on a single time point (30min after α-IgM treatment), we investigated the temporal dynamics of MEK and ERK phosphorylation in BL-2 cells (**FIGURE 3A**). An increase in MEK and ERK phosphorylation as early as 2 min after α-IgM-mediated BCR stimulation is observed in the presence of the p38 inhibitor SB203580. This p38 inhibitor-dependent increase of MEK/ERK phosphorylation persists for the entire span of the measured 60 minutes post stimulation with the strongest induction within the first 10 minutes and a slight decrease thereafter (**FIGURE 3A**).

**FIGURE 3.**
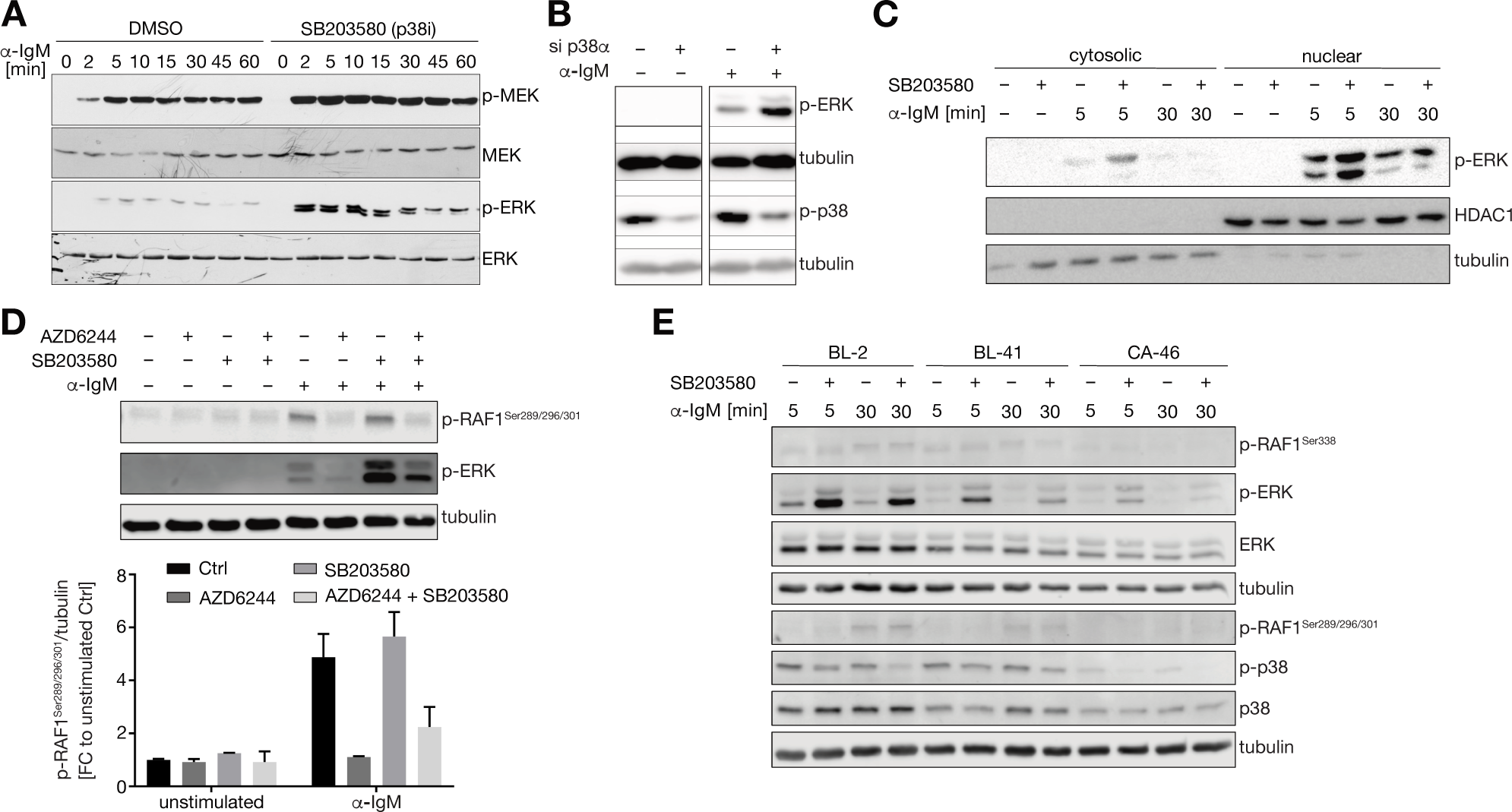
Increased MEK/ERK-pathway activity in B cell receptor activated B-cell lines after p38 intervention. (A) Changes in the phosphorylation of MEK and ERK in BL-2 cells after treatment with α-IgM in the presence or absence of the p38 inhibitor SB203580. (B) Phosphorylation of ERK is further increased in α-IgM-treated BL-2 cells after 24h of p38α (MAPK14) knockdown. (C) Phosphorylation of ERK is enriched within the nucleus of α-IgM treated BL-2 which is further enhanced by inhibiting p38. Tubulin and HDAC1 were used as reference for the cytosolic and nuclear fraction, respectively. (D) (TOP) Phosphorylation of c-RAF at serine-residues 289/296/301 is increased after 30 min α-IgM treatment in BL-41 cells but not affected by p38 inhibition. The inhibition of MEK using AZD6244 interrupts the phosphorylation of RAF. Representative Western blot. (BOTTOM) Bar plots quantifying c-RAF phosphorylation measurements for n=2 replicates. (E) p38 affects the MEK/ERK pathway in a comparable way in different BL cell lines after α-IgM treatment. Shown are phosphorylations of Raf1 at serine 338 (activatory site) and 289/296/301 (ERK feedback sites) as well as of ERK and p38 in Burkitt lymphoma cells BL-2, BL-41 and CA-46.

To directly asses the involvement of p38 in this predicted cross talk, a knockdown of p38α (MAPK14) was performed. P38α is one of the most abundantly expressed isoforms of p38 in BL-2 cells (**Suppl. Figure 2**). Importantly, the knockdown of p38α affects the phosphorylation of ERK similarly to the previous p38 inhibition (**FIGURE 3B**). This further supports the observation that p38 activity dampens the MEK/ERK pathway and that in BL-2 cells this seems to be conferred by the α-isoform of p38.

By analyzing subcellularly fractionated cell lysates, it can be shown that both cytosolic as well as nuclear pERK is increased in the presence of the p38 inhibitor SB203580 in α-IgM-stimulated BL-2 cells (**FIGURE 3C**). As only active ERK is able to enter the nucleus but not MEK (Witzel et al., 2012) it is unlikely that the cause of the upregulation is a prolonged retention of ERK together with active MEK in the cytosolic scaffold. This finding also indicates that ERKs activity is likely to be propagated to its many nuclear targets even in the presence of p38 inhibitors.

Having verified the negative crosstalk from p38 on MEK and ERK, the next step was to characterize the molecular mechanism that underlies this crosstalk. The RAF-MEK-ERK signaling cascade is a well-characterized MAPK pathway involved in different cellular processes initiated by different cell surface receptors and RAS. Upstream kinase RAF is known to be negatively regulated by phosphorylation at serines 289, 296 and 301, which are known negative feedback sites phosphorylated by ERK (Dougherty et al., 2005). Since p38, as ERK, belongs to the family of MAPK we examined whether p38 regulates phosphorylation at serine 289, 296 and 301 and thus might affect RAF1 activity. After 30 minutes of α-IgM stimulation, RAF1^S289/296/301^ phosphorylation is clearly detectable and diminished by MEK inhibition using AZD6244, confirming the regulation by ERK also within the BCR signaling cascade (**FIGURE 3D**). However, RAF1 phosphorylation at these sites is not affected by p38 inhibitor SB203580 (**FIGURE 3D**). Also, in BL-2 and CA-46 Burkitt lymphoma cells no difference on the RAF1 feedback sites can be seen (**FIGURE 3E**). This indicates that p38 must attenuate the MEK-ERK pathway directly after BCR activation while the feedback from ERK to RAF1 seems p38-independent.

Next to the feedback we also checked upstream activation of RAF1 by measuring phosphorylation at serine 388 (Diaz et al., 1997) in three different BL cell lines BL-2, BL-41 and CA-46 (**FIGURE 3F**). Phosphorylation of S338 increases slightly after BCR activation, however, there is no effect by p38 inhibition (**FIGURE 3E**). Importantly, in all three cell lines p38 inhibition affects the MEK/ERK pathway in a comparable way after α-IgM treatment as shown by upregulation of pERK (**FIGURE 3E**). Therefore, our data support the view that this crosstalk is a widely existing phenomenon in BL cells. In addition, we also saw that CD40L stimulation in CA-46 cells could also produce hyperactivation of pERK together with p38 inhibition (**Suppl. FIGURE 3**). This demonstrates that the observed p38-ERK crosstalk is not only limited to BCR activation, but is an inherent mechanism in BL signaling.

### Global phosphoproteomic analysis of BL-2 cells supports model-derived pathway network crosstalk

To further characterize the crosstalk from p38 to MEK/ERK and to answer the question about a more general consequences on overall signaling, a systematic proteomic analysis was performed. Specifically, we performed phosphoproteomic mass spectrometry combined with the tandem mass tag (TMT) technology that allows to quantitatively compare phosphorylation in different samples without drop outs (Mertins et al., 2018).

Phosphoproteomes were obtained for unstimulated controls, and cells stimulated for 30 min with α-IgM that were incubated with inhibitors of p38 (SB203580), PI3K (BKM120) and mTORC1 (Rapamycin) or solvent control (DMSO) as in the above-described analysis. PI3K inhibitor was chosen as it represents a central hub in the deduced network model with mTORC1 the target of rapamycin being a major downstream mediator, whereas the p38 inhibition was included to get an unbiased insight into the downstream effects of p38 in BCR signaling as well as to extend the understanding of the p38-MEK/ERK crosstalk in B cells.

Overall, 28871 phosphosites from 5698 proteins (localization probability>0.75) were identified reliably. When investigating the global effect on the 3000 most varying phosphosites by hierarchical clustering (**FIGURE 4A**) and principal component analysis (**FIGURE 4B**), it is evident that α-IgM treatment had the strongest impact. Interestingly, inhibition of PI3K by BKM120 reverted the effect of α-IgM partially, and thus α-IgM+PI3Ki samples were located between Control+DMSO and α-IgM+DMSO samples in PC1-PC2 space. α-IgM treatment in combination with Rapamycin (mTORC1) or SB203580 (p38) clustered and colocalized well with α-IgM+DMSO treatment. This indicates that the inhibition of mTORC1 and p38 had a rather confined influence on the phosphoproteome in α-IgM activated BL-2 cells.

**Figure 4:**
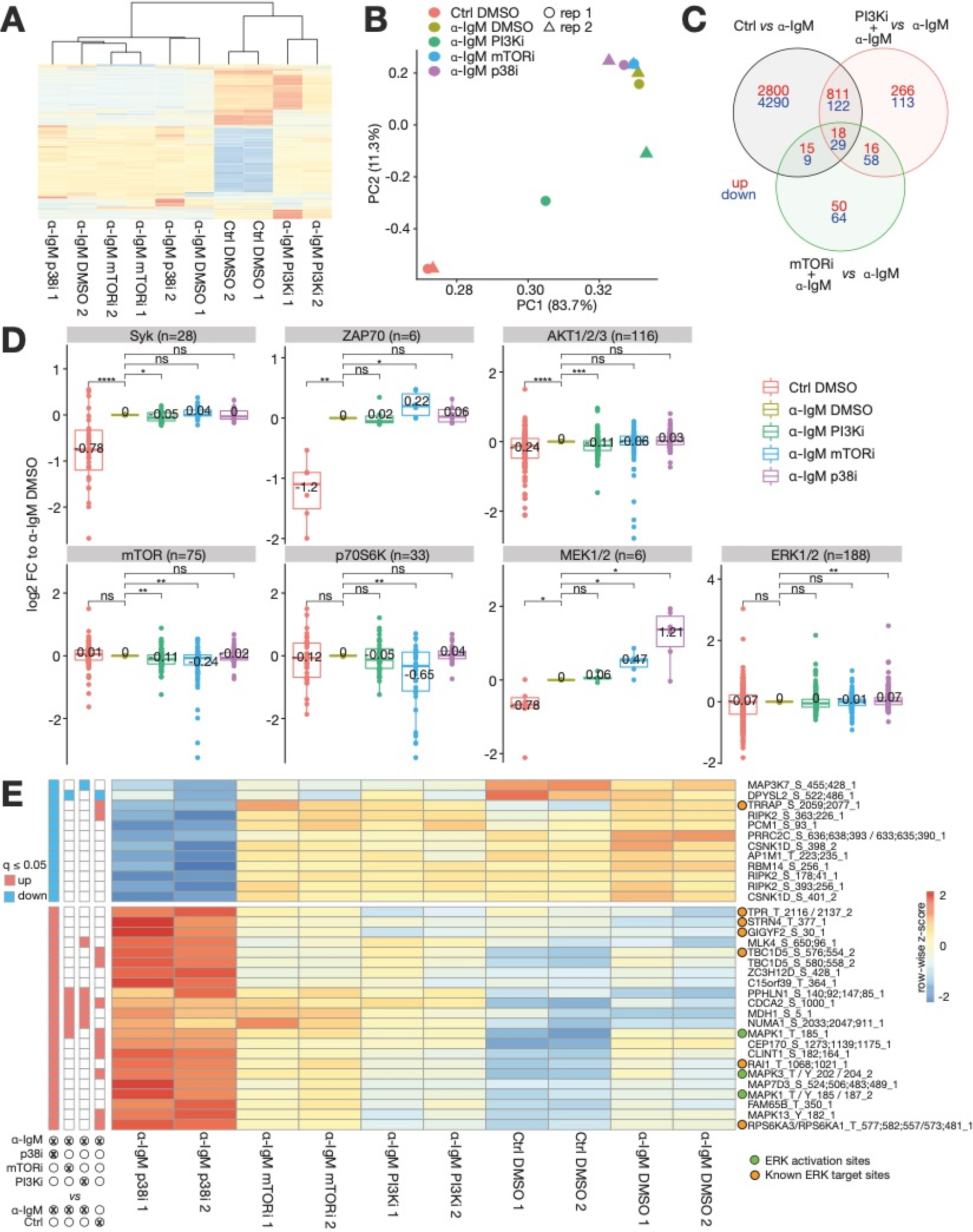
Phosphoproteomics analysis supports the BCR-signaling model and reveals a dominant effect of the PI3K pathway inhibition onto BCR-signaling in BL-2 cells. Analysis of Tandem-Mass-Tag (TMT) Mass spectrometry measurements for BL-2 cells treated with α-IgM and inhibitor solvent DMSO or inhibitors of PI3K (BKM120), MTORC1 (Rapamycin) or p38 (SB203580) (n=2). (A) Hierarchical clustering of 3000 most varying phosphosites demonstrates a global effect of α-IgM and PI3K inhibitor BKM120 on the phosphoproteome and subtle effects of the remaining inhibitors. (B) Principal component analysis shows that α-IgM effect is governing the principal component 1 and 2. Only PI3Ki treatment is able to partly revert the α-IgM effect. (C) Overlap of differentially regulated phosphosites (limma, FDR≤5%) for indicated selected comparisons. (D) Upstream kinase activity assessment on base of log2 fold changes (vs. α-IgM+DMSO) in PhosphoSitePlus-annotated target sites (Nov 2021) for selected kinases. Significance asserted by two-sided t-test: ns - not significant; * - <u>0.05</u>; ** - <u>0.01</u>; *** - <u>0.001</u>; **** - 0.0001); Average value indicated. (**E**). Phosphosites significantly regulated by p38 inhibitor SB203580 (limma, FDR≤5%). Left panel denotes which site was found to be significantly regulated (blue - down; red – up) by the indicated comparison. Sites are annotated as follows: ‘*HGNC symbol’*_’*amino acid’*_’*position’*_’*number. of phosphosites’*; ERK activation sites and known target sites of ERK (Ünal et al., 2017) are indicated by green and orange circles, respectively.

Differential expression analysis showed that α-IgM treatment had the largest effect with 8094 phosphosites (28% of detectable phosphosites) that were significantly different compared to untreated control (limma FDR<5%, **FIGURE 4C**).). PI3K inhibition had an influence on 1433 phosphosites when compared to α-IgM treatment alone. mTORC1 inhibition led to a significant change in only 259 phosphosites, of which 47% are shared sites with its upstream regulator PI3K. With 12%, a considerable part of the α-IgM-regulated phosphoproteome is counter-regulated by PI3K (980/8094). Interestingly this PI3Ki counter-regulation is much more prominent for α-IgM down-regulated sites (23%) than for α-IgM upregulated sites (3%). Due to its large counter-active potential, it can be assumed that PI3K is a very potent target for treating BCR-addicted neoplasms.

Differences in substrate phosphorylation are usually caused by differences in kinase activity. Therefore, upstream kinase annotation provided by the PhosphoSitePlus Database (Hornbeck et al., 2015) was used to get an insight into the potential kinase activity by a combined analysis of their target sites (**FIGURE 4D**). Significant kinase activity changes were defined by testing all treatments to α-IgM treatment alone using a pairwise t-test. Kinases acting further upstream such as Syk and ZAP70 are most strongly regulated by α-IgM vs Control treatment. The effect of inhibitors on those upstream kinases was rather small with marginally significant effects of PI3Ki counteracting SYK-target induction and mTORi increasing ZAP70 target induction (**FIGURE 4D**). Reassuringly, *bona fide* target kinases for the inhibitors, i.e., AKT for PI3K and mTOR/p70S6K for mTOR showed the strongest and most significant reversal of α-IgM induction with their respective inhibitor treatment. Importantly, the proposed p38-MEK/ERK crosstalk is also visible in the significant upregulation of MEK and ERK target site phosphorylations after SB203580 treatment. This further indicates that the crosstalk is functional and inhibition of p38 leads to an enhanced ERK signaling output on a global scale.

Having ascertained the negative p38-MEK/ERK crosstalk, the significantly regulated sites (limma, FDR<=5%) by SB203580 were further investigated (**FIGURE 4E**). p38 inhibitor responsive phosphosites are rather specifically regulated, with 70% (24/34) of significantly regulated sites neither co-regulated by other inhibitors nor counteractively regulated by α-IgM treatment. 12 sites are strongly downregulated and 22 hyperactivated compared to Control and DMSO treatment. Three of the hyperactivated sites are activatory phosphosites of ERK1/2 (red) demonstrating a direct ERK hyperactivation. We then checked other phosphosites for known ERK-targets, using a previously compiled compendium of ERK targets (Ünal et al., 2017). Seven of the p38 inhibitor-dependent phosphosites are described as ERK targets, of which six follow the same hyperactivation pattern as ERK phosphorylation itself. This further corroborates that next to ERK also downstream partners are hyperactivated by p38 inhibition. Notably TRRAP, the only identified known ERK target to be downregulated by p38 inhibition seems to be a shared target of both ERK and p38. This can be proposed based on a SILAC-based phosphoproteomics study which observed, that TRRAP^S2077^ is upregulated after 15 min EGF stimulation and downregulated by EGF stimulation and p38 inhibitor (SB202190) treatment in HeLa cells (Pan et al., 2009). Taking into account the similarity of the results in that study and our own observation we conclude that TRRAP^S2077^ is dominated by p38 over ERK activity.

From the phosphoproteome analysis, we conclude that p38 inhibition indeed leads to stronger ERK activation, but it is rather confined to a small subset of the signaling network. While the mechanism of the crosstalk remains unclear, the phosphoproteome does identify 19 candidate phosphosites that are p38-regulated but not known ERK targets which could be investigated in future studies.

### BL-2-dervied network improves starting basis for network development of DLBCLs

Having developed a semi-quantitative model for activated BCR signaling in BL cells we sought to investigate if a similar signaling network exists in cells with aberrant BCR signaling such as in ABC-like DLBCL. For that reason, the intracellular signaling networks of two representative cell lines HBL-1 and OCI-LY3 were analyzed. Similarly, to BL-cell lines earlier (cf. **FIGURE 1A**), the signaling in both cell lines was perturbed using the same inhibition and measuring scheme. Additional α-IgM stimulation was not required, as both cell lines exhibit chronic active BCR signaling (Davis et al., 2010). The resulting phosphoprotein perturbation data of both cell lines (**FIGURE 5A**) were analyzed using our STASNet pipeline.

**Figure 5:**
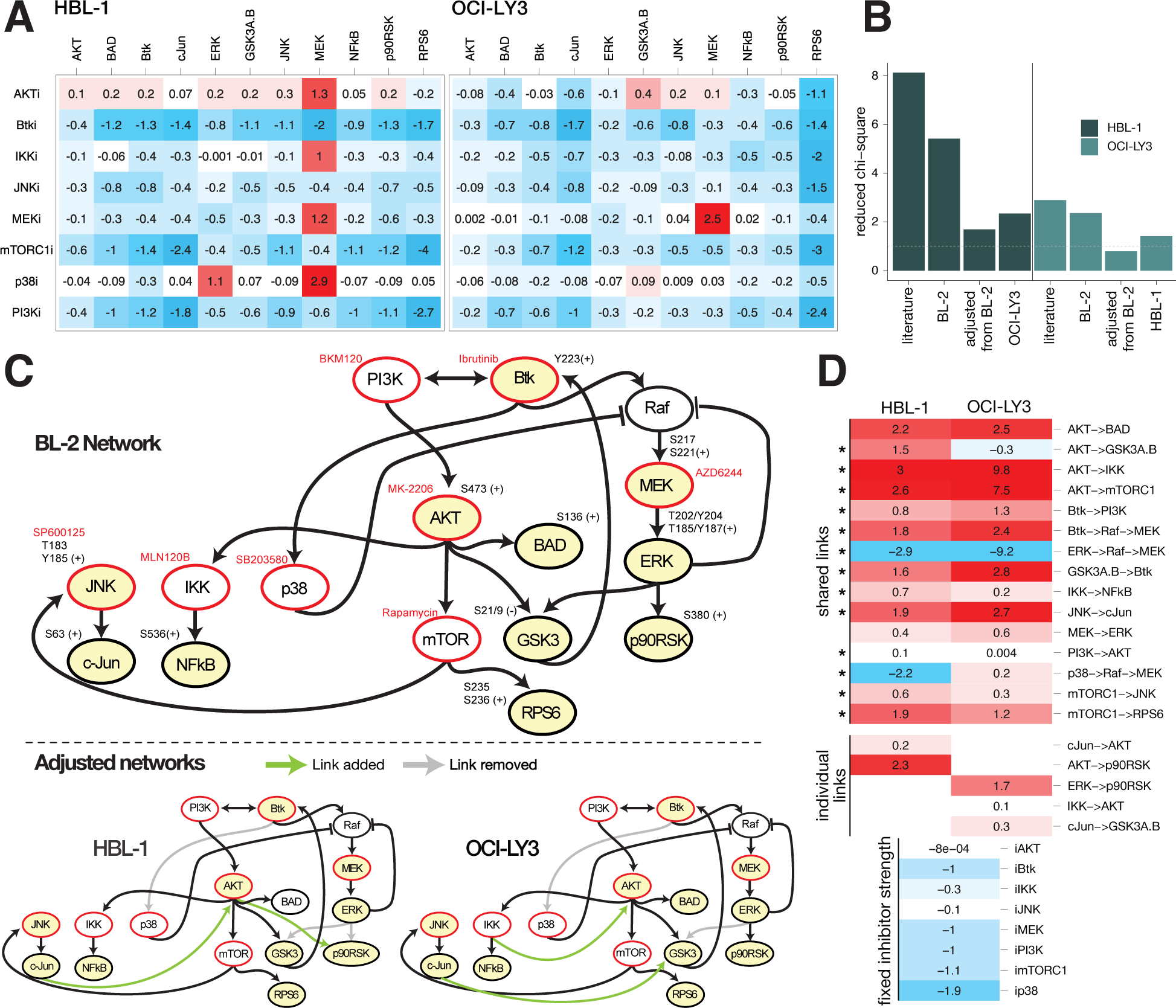
BL-2-derived network structure sets a veritable starting base to develop networks for DLBCL cell lines HBL-1 and OCI-LY3. (**A**) Systematic perturbation data of MagPix measurements of the DLBCL cell lines HBL-1 and OCI-LY3 quantified as log2 fold changes to solvent control (DMSO); mean of n=3. (**B**) Goodness of fit expressed as reduced chi-square statistic (Xr) on selected network structures for the two DLBCL cell lines HBL-1 and OCI-LY3. *literature* – network from **Figure 1B**; *BL-2* – BL-2 network derived from **Figure 1C**; *adjusted from BL-2* – BL-2 network locally adjusted to respective DLBCL cell line; *OCI-LY3/HBL-1* – final adjusted network of respective other DLBCL cell line (see **C**). (**C**) Network structures of BL-2 derived starting network and final DLBC-specific networks trained on HBL-1 and OCI-LY3 data (see **A**). (**D**) Side-by-side comparison of the model pathway coefficients (log scale) with fixed inhibitor strengths set to mean of both cell line models. Empty tiles indicate missing links in one of the cell lines (individual links). Asterisks point to non-overlapping confidence intervals as estimated by STASNet profile likelihood function.

We first tested if the literature-based network or the BL-2-derived network provides a better starting point for the development of the signaling model. We trained models for both cell lines on (a) the literature network (cf. **FIGURE 1D** grey and black arrows) and (b) BL-2-derived final network (cf. **FIGURE 1D**) without receptor proximal signaling as no stimulation perturbation was present. For both DLBCL cell lines, we found that the BL-2-derived model resulted in a better fit compared to the literature-based model (**FIGURE 5B**). We then structurally adjusted the BL2-network to the respective DLBCL-cell line reaching reduced chi-square residuals close to the theoretical expectance for optimal fit, i.e., 1 (**FIGURE 5B** – adjusted from BL-2). We also tested whether those cell line-specific fits yield generality in DLBCLs and re-fitted HBL-1 and OCI-LY3 data with the best network structure identified for the respective other cell line. Since both cross-fittings resulted in clearly higher residuals than the respective self-best network fit, we decided to use the individually developed networks for the further analysis.

The corresponding network starting from the BL-2 BCR pathway model is presented in **FIGURE 5C** before and after training on HBL-1 and OCI-LY3 cell lines. In both cell lines the links from BTK to p38 and ERK to GSK3 were removed by the network adjustment strategy of STASNet (**FIGURE 5C**). In HBL-1 cells additional links from JNK/cJun to AKT and from AKT to p90RSK were required, whereas in OCI-LY3 the network required a link between JNK/cJun and GSK3 and between IKK and AKT for optimal fits. The link from ERK to p90RSK was removed only in the HBL-1 network. In agreement with the findings from the analysis of the Burkitt lymphoma cell line BL-2, we also note a central role of PI3K in the studied DLBCLs indicated by the network central position of the PI3K node even after the individual network adjustment (**FIGURE 5C, bottom**), i.e., every network node can be reached by PI3K. In addition, when inspecting the links removed and added in both DLBCL cell lines, a shift from less control by MAPK to more control by mTOR/AKT can be noticed which further corroborated the importance of PI3K in DLBCL signaling. This PI3K/mTOR-dependency is further underlined by the growth inhibitory effect of corresponding inhibitors in those cell lines (see **Suppl. Figure 4**).

Next to identifying the structural similarities and differences, STASNet allows to reveal the quantitative signalling differences in the cell lines by comparing the pathway coefficients (**FIGURE 5D**). For better comparability we fixed the inhibitor strength to the mean of both cell line models. After applying parameter stability analysis using profile likelihood, we noted that of the 15 shared links only 2 (AKT->BAD, MEK->ERK) are not significantly different (i.e., 95% confidence intervals do not overlap; **Suppl. Table 4**). Of note the p38->Raf->MEK crosstalk is negative in HBL-1-like in BL-2 and BL-41 - but faintly positive in OCI-LY3.

We found that from 18 identifiable pathway coefficients that are present in the BL-2 network model aside from receptor-proximal signaling (**FIGURE 5C TOP**), 15 (83%) and 16 (89%) coefficients were also required in HBL-1 and OCI-LY3 cell line models, respectively. Next to the removal of three (HBL-1) and two (OCI-LY3) links, both models required only two additional pathway coefficients for their final best fit, suggesting that many insights from α-IgM-stimulated BL cells can be transferred to DLBCLs. However, these changes are necessary, as we observe a ∼3-fold improvement of fit, i.e., reduction of Xr, in both DLBCL cell lines with five and four adjustments for HBL-1 and OCI-LY3, respectively (**FIGURE 5B**). This indicates that a common model of BL and DLBCL would perform poorly and local adjustments are still required but that a common conserved core network of BCR downstream signaling exists. Altogether this study demonstrates that a conserved BCR network found by studying acute BCR signaling (BL cell lines) was largely transferable to study chronic BCR signaling (DLBLC cell lines) indicating that the aberrant activation did not lead to a major signaling rewiring.

## Conclusion

In this work, we established a (semi)-quantitative signaling network of the BCR and tested it in different B cell lymphoma cell lines. We explored how a more quantitative understanding of signaling can be used to describe a conserved B cell receptor regulated signaling network, which can be used as a starting point to quantify cell-specific, individual networks. The development of the new BCR network model allowed us to introduce so far not described links in BCR mediated intracellular signaling. Earlier described network cross talks and feedbacks were confirmed, but also new feedbacks are identified further supporting the complexity of the BCR signaling network. Thus, the well-known RAF/MEK/ERK regulatory loop also exists in BLs and DLBCLs, whereas the negative impact of BCR mediated p38 activation onto this loop is a newly discovered crosstalk found in three BL cell lines BL-2, BL-41 and CA-46 and one DLBCL cell line HBL-1. We propose that the identified negative crosstalk from BCR mediated activation of p38 to the RAF/MEK/ERK pathway might dampen an excessive RAF/MEK/ERK signaling cascade activated by α-IgM stimulation in BL or present in individual DLBCL. Despite the discovery of feedbacks and crosstalks in ERK and p38 signalling we could not find a cellular dependency in the cell lines we studied. However, as we determined that those signaling motifs are conserved across different B cell lymphoma types it might be possible that insights of this study apply also to other B cell lymphomas. For example, a recent study described an essential dependency to MAPK activity in Chronic Lymphocytic Leukemia progression suggesting that insights from this study could be valuable for treatment decisions in CLL (Ecker et al., 2023).

While the mechanism behind the p38 – MEK/ERK crosstalk remains enigmatic, it is clear that it is neither mediated by the ERK feedback sites nor the activatory site of RAF-1. A previous study on Hela cells determined that the crosstalk is conferred by direct binding of p38 to ERK (Zhang et al., 2001), as it demonstrated direct binding and no change in MEK phosphorylation. In our study we confirmed that MEK phosphorylation effected in cells that show the p38 crosstalk when treated with p38 inhibitor (**FIGURE 2F, 3A, 5A**). This upregulation is incompatible with the negative feedbacks downstream of ERK as this would result in an attenuation of pMEK following the observed ERK hyperactivation. Therefore, we postulate that crosstalk acts on or even more probable upstream of MEK.

In addition, a positive link between MTOR and JNK and a positive feedback downstream of AKT to upstream of PI3K/Btk is proposed in our consensus model, which notably are kept also in both DLBCL cell line models. Importantly, most observed pathway crosstalks are part of the established core BCR network, whereas other are more cell type and cell line specific. This supports the view that our model reflects common elements of intracellular signaling of α-IgM stimulated BL cell lines.

In conclusion, we could demonstrate that the more generic network trained from BL-2 and BL-41 cell lines had better transfer capacity to DLBCL cell lines HBL-1 and OCILY3 than the literature network indicating that a model trained on BCR-activated B-cells is informative to be transferred to BCR-driven cancer models. Importantly, our conserved BCR network model can be applied to the situation in antigen-driven BCR activation (BL-2/BL-41) and the situation, were the BCR pathway is antigen independently activated, driven by gain-of-function mutations in specific signaling molecules of the BCR pathway (HBL-1/OCI-LY3). A recent study that compared primary Chronic Lymphocytic Leukemia and non-malignant B Cell signaling by modeling a smaller network (5 nodes) using Bayesian inference learning underscores our finding by determining only a modest topology difference between the malignant and non-malignant network (Wolf et al., 2022). In here we can recapitulate this finding on a larger network for aberrant and non-aberrant B-cell receptor driven signaling.

## Acknowledgments

We thank all members of the joint project MMML-Demonstrator for helpful discussion in running the analysis. This work was funded by grants from the Federal Ministry of Education and Research (BMBF) within the joint project e:Med MMML-Demonstrator (031A428B and 031A428G). BK was funded by Deutsche Krebshilfe (70114307), DKFZ (Young Investigator Grant) and Bundesinstitut für Risikobewertung (60-0102-01.P61).

## Author contributions

I.R., H.K., V.K. and A.S. performed the experimental work. MLK and PM did the TMT phosphoproteome analysis, B.K. did the data analyses and interpretation. B.K., A.K., N.B. and D.K. designed the research, analyzed and interpreted data, and wrote the manuscript.

## Data and materials availability

Mass spec data are available via ProteomeXchange with identifier PXD047709. All relevant data and pipelines for modelling can be accessed on the zenodo repository https://zenodo.org/doi/10.5281/zenodo.10732059.

## Supplementary Materials

**Supplemental Figure 1.**
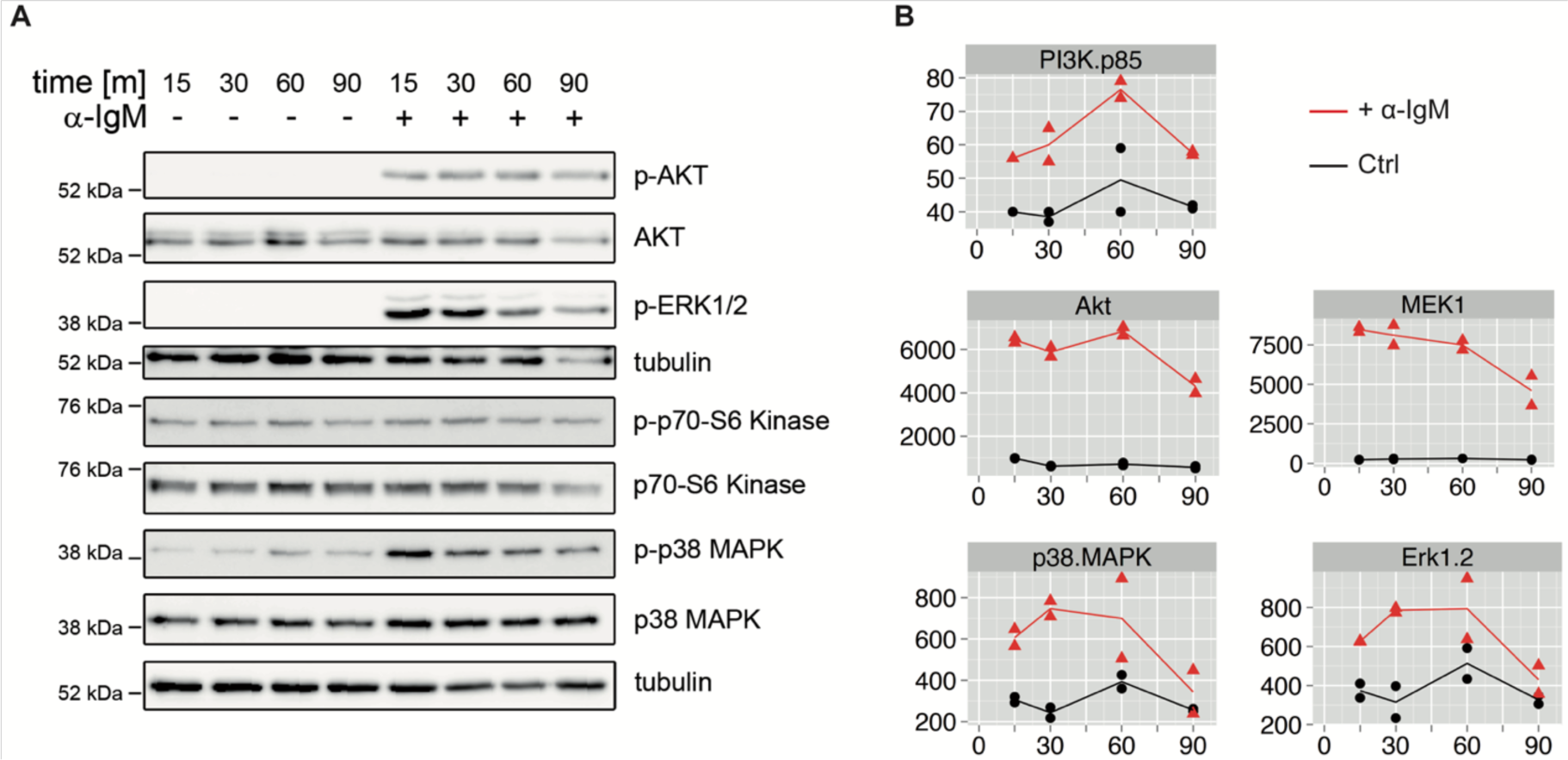
Changes in the phosphorylation of AKT, ERK1/2, p70S6K and p38/MAPK14 over time after treatment of BL-2 cells with α-IgM. The phosphorylation changes were estimated by (A) immunoblot and (B) Bead-based ELISA analysis on the same samples depicted in fluorescence intensities.

**Supplementary Figure 2.**
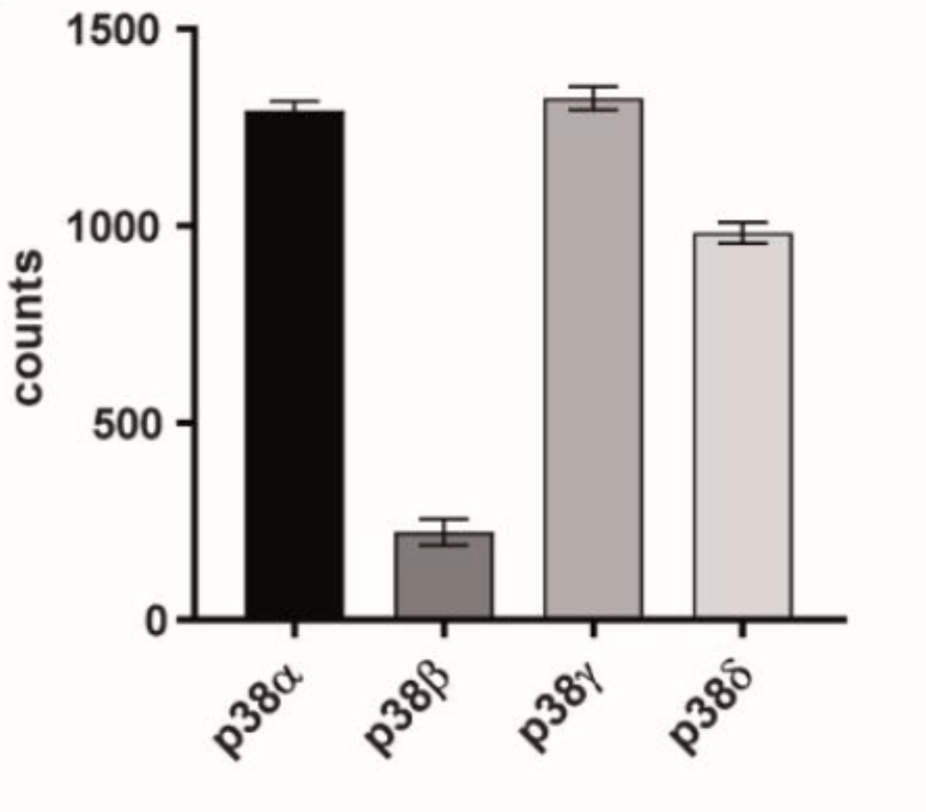
p38α/MAPK14 is one of the most abundantly expressed isoforms of p38 in BL-2. Total reads of the p38 subunits α (MAPK14), β (MAPK11), γ (MAPK12), δ (MAPK13) are displayed from RNA sequencing analysis of BL-2 cells (n=3, data from microarrays conducted at Wolff et al., 2018).

**Supplemental Figure 3.**
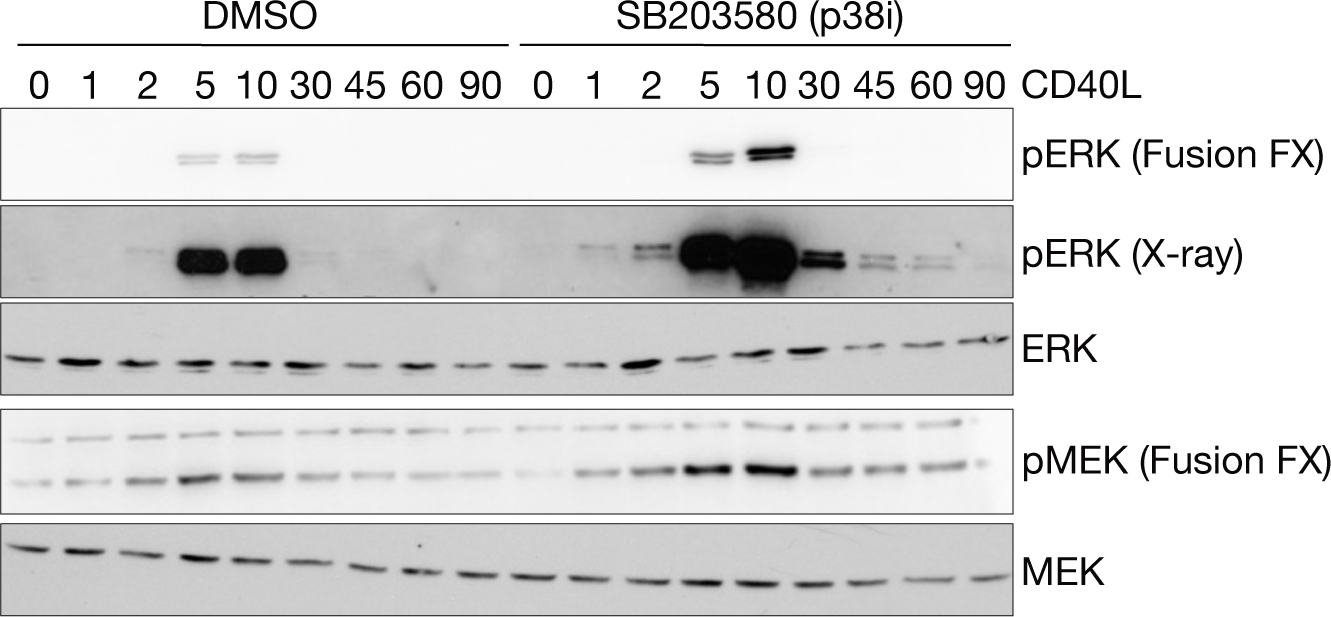
Increased MEK/ERK-pathway activity after p38 MAPK14 intervention can be observed also in response to CD40-Ligand treatment in BL cells. CA-46 BL cells were treated with CD40 ligand for up to 90 min without or with 2µM p38 inhibitor SB203580. Data were analyzed by classical IB-chemoluminescence imaging and by the Fusion-FX platform.

**Supplemental Figure 4.**
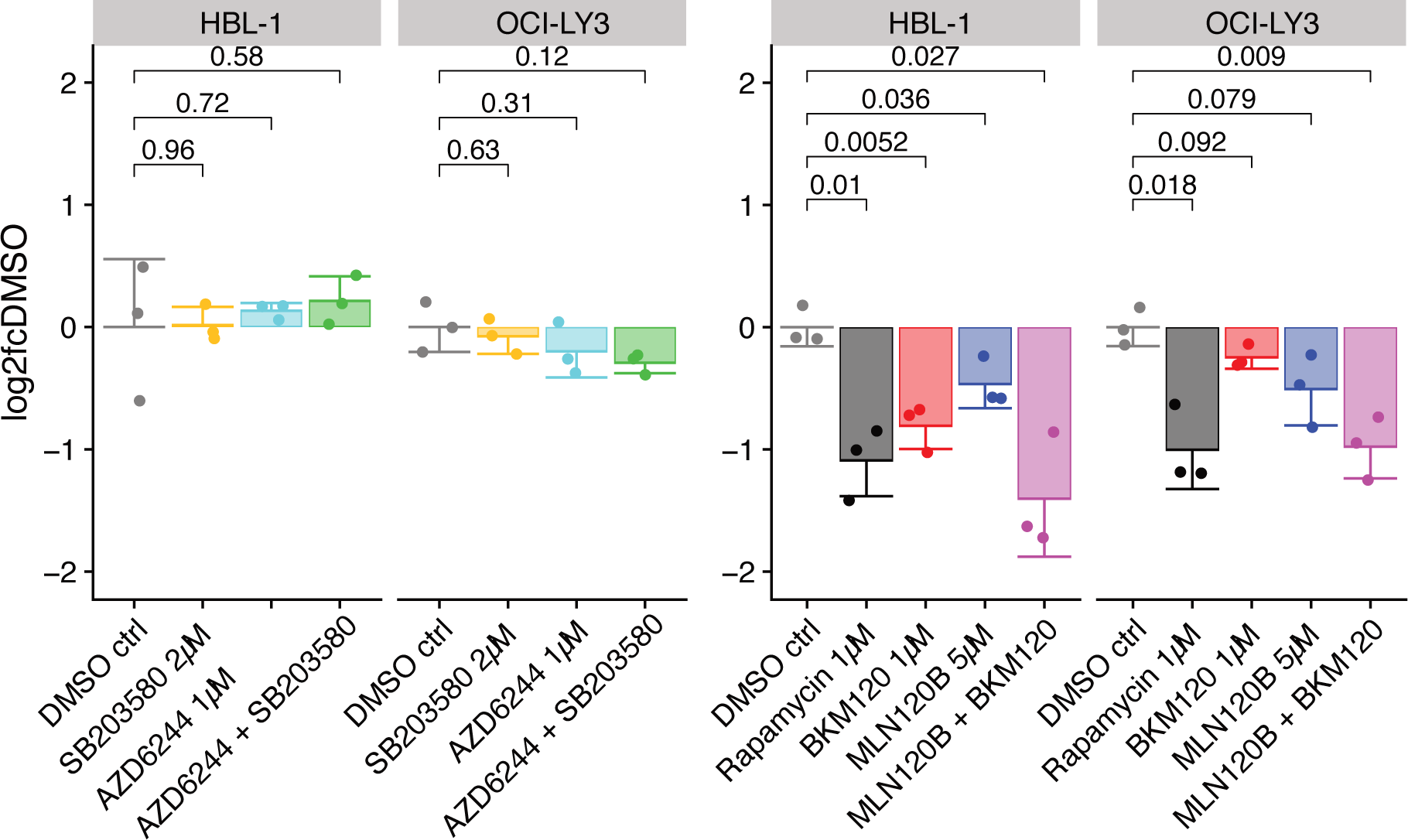
Relative growth of DLBCL cells after drug treatments. Cell counts of indicated DLBCL cell lines treated for 72hwith indicated inhibitors and concentrations, as log2 fold change to respective mean solvent control (DMSO). P-values derived by two-sided T-test; n=3. Inhibitors (target) : SB203580 (p38), AZD6244 (MEK), Rapamycin (mTORC1), BKM120 ( PI3K) and MLN120B (IKK).

**Supplemental Table 1.** Side-by-side model coefficients and confidence intervals for BL-2 and BL-41 data with fixed inhibitor quantifications. Parametrization for the best fit and in brackets the upper and lower boundaries of the 95% confidence interval, derived by profile likelihood (alpha = 0.05, 1 degree of freedom). ni denotes nonidentifiable confidence intervals in that direction (i.e., alteration of the parameter can be compensated by changing other model parameters). Model coefficients with non-overlapping confidence intervals are termed significantly different between BL-2 and BL-41.

| Model coefficients | BL-2 | BL-41 | Significant difference |
| --- | --- | --- | --- |
| r_BAD_AKT | 0.69 (0.4 – 1.24) | -0.08 (-0.22 – 0.06) | YES |
| r_GSK3A.B_AKT | 0.37 (0.18 – 0.74) | -0.5 (-0.58 – -0.43) | YES |
| r_IKK_AKT | 2.52 (0.09 – ni) | 0.12 (0.04 – 0.19) | NO |
| r_mTORC1_AKT | 2.63 (0.12 – ni) | -0.49 (-0.59 – -0.42) | YES |
| r_BCR_α-IgM*r_Syk_BCR | 0.52 (0.43 – 0.6) | 0.64 (0.54 – 0.72) | NO |
| r_PI3K_Btk | 5.36 (0.67 – ni) | 19.83 (17.39 – 23.1) | NO |
| r_MEK_Raf*r_Raf_Btk | 107.52 (76 – 233) | 169.28 (159 – 180) | NO |
| r_MEK_Raf*r_p38_Btk*r_Raf_p38 | -32.86 (-43 – -22) | -75.93 (-85 – -65) | YES |
| r_GSK3A.B_ERK | 0.27 (0.11 – 0.44) | 0.0019 (-0.11 – 0.1) | YES |
| r_MEK_Raf*r_Raf_ERK | -9.13 (-15.7 – -4.8) | -9.17 (-14.5 – -7.6) | NO |
| r_p90RSK_ERK | 0.36 (0.16 – 0.57) | 0.22 (-0.24 – 0.67) | NO |
| r_ZAP70_GSK3A.B | 1.2 (0.71 – 2.13) | 1.54 (1.26 – 1.81) | NO |
| r_NFkB_IKK | 0.22 (0.08 – ni) | -1.01 (-1.62 – -0.42) | YES |
| r_cJun_JNK | 0.91 (0.57 – 2.43) | 25.81 (3.59 – ni) | YES |
| r_ERK_MEK | 0.3 (0.23 – 0.36) | 0.18 (0.15 – 0.21) | YES |
| r_AKT_PI3K | 1.58 (0.45 – ni) | -1.17 (-1.41 – -1.02) | YES |
| r_Btk_Syk | 0.1 (0.02 – 0.14) | 0.05 (0.03 – 0.06) | NO |
| r_PI3K_Syk | -0.56 (-2.29 – -0.05) | -0.21 (-0.42 – -0.01) | NO |
| r_Btk_ZAP70 | 0.12 (0.07 – 0.19) | 0.03 (0.01 – 0.03) | YES |
| r_JNK_mTORC1 | 0.35 (0.1 – ni) | 0.04 (0.03 – 0.04) | YES |
| r_RPS6_mTORC1 | 0.31 (0.08 – ni) | 0.8 (0.63 – 0.97) | NO |

**Supplemental Table 2.** Side-by side model coefficients and confidence intervals for HBL-1 and OCI-LY3 data with fixed inhibitor quantifications. Parametrization for the best fit and in brackets the upper and lower boundaries of the 95% confidence interval, derived by profile likelihood (alpha = 0.05, 1 degree of freedom). ni denotes nonidentifiable confidence intervals in that direction (i.e., alteration of the parameter can be compensated by changing other model parameters). Model coefficients with non-overlapping confidence intervals are termed significantly different between HBL-1 and OCI-LY3. – indicates missing link in either cell line.

| Model coefficients | HBL-1 | OCI-LY3 | Significant difference |
| --- | --- | --- | --- |
| r_BAD_AKT | 2.23 (1.78 – 2.67) | 2.5 (1.61 – 3.42) | NO |
| r_GSK3A.B_AKT | 1.51 (1.31 – 1.67) | -0.27 (-0.27 – -0.2) | YES |
| r_IKK_AKT | 3.02 (2.35 – 3.68) | 9.8 (9.39 – 10.04) | YES |
| r_mTORC1_AKT | 2.62 (2.38 – 2.91) | 7.51 (7.13 – 7.88) | YES |
| r_p90RSK_AKT | 2.34 (1.89 – 2.78) | – | – |
| r_PI3K_Btk | 0.84 (0.72 – 0.96) | 1.3 (1.3 – 1.36) | YES |
| r_MEK_Raf*r_Raf_Btk | 1.83 (1.27 – 2.39) | 2.39 (2.39 – 5.3) | YES |
| r_MEK_Raf*r_Raf_ERK | -2.92 (-4.02 – -2.14) | -9.24 (ni – -5.71) | YES |
| r_Btk_GSK3A.B | 1.59 (1.36 – 1.8) | 2.8 (2.5 – 2.94) | YES |
| r_NFkB_IKK | 0.67 (0.52 – 0.81) | 0.22 (0.13 – 0.31) | YES |
| r_cJun_JNK | 1.89 (1.66 – 2.11) | 2.71 (2.48 – 2.83) | YES |
| r_ERK_MEK | 0.44 (0.31 – 0.62) | 0.6 (0.39 – 0.6) | NO |
| r_AKT_PI3K | 0.14 (0.11 – 0.17) | 0.0036 (0.0036 – 0.02) | YES |
| r_MEK_Raf*r_Raf_p38 | -2.22 (-2.74 – -1.71) | 0.16 (-0.43 – 0.86) | YES |
| r_AKT_cJun | 0.15 (0.12 – 0.18) | – | – |
| r_JNK_mTORC1 | 0.64 (0.57 – 0.7) | 0.25 (0.25 – 0.26) | YES |
| r_RPS6_mTORC1 | 1.9 (1.75 – 2.04) | 1.25 (1 – 1.57) | YES |
| r_p90RSK_ERK | – | 1.65 (0.99 – 2.24) | – |
| r_AKT_IKK | – | 0.1 (0.1 – 0.11) | – |
| r_GSK3A.B_cJun | – | 0.28 (0.28 – 0.29) | – |

